# Dual mTORC1/mTORC2 inhibition as a Host-Directed Therapeutic Target in Pathologically Distinct Mouse Models of Tuberculosis

**DOI:** 10.1101/2021.02.10.430715

**Authors:** Rokeya Tasneen, Deborah S. Mortensen, Paul J. Converse, Michael E. Urbanowski, Anna Upton, Nader Fotouhi, Eric Nuermberger, Natalie Hawryluk

## Abstract

Efforts to develop more effective and shorter-course therapies for tuberculosis have included a focus on host-directed therapy (HDT). The goal of HDT is to modulate the host response to infection, thereby improving immune defenses to reduce the duration of antibacterial therapy and/or the amount of lung damage. As a mediator of innate and adaptive immune responses involved in eliminating intracellular pathogens, autophagy is a potential target for HDT in tuberculosis. Because *Mycobacterium tuberculosis* modulates mammalian target of rapamycin (mTOR) signaling to impede autophagy, pharmacologic mTOR inhibition could provide effective HDT. mTOR exists within two distinct multiprotein complexes, mTOR complex-1 (mTORC1) and mTOR complex-2 (mTORC2). Rapamycin and its analogs only partially inhibit mTORC1. We hypothesized that novel mTOR kinase inhibitors blocking both complexes would have expanded therapeutic potential. We compared the effects of two mTOR inhibitors: rapamycin and the orally available mTOR kinase domain inhibitor CC214-2, which blocks both mTORC1 and mTORC2, as adjunctive therapies against murine TB, when added to the first-line regimen (RHZE) or the novel bedaquiline-pretomanid-linezolid (BPaL) regimen. Neither mTOR inhibitor affected lung CFU counts after 4-8 weeks of treatment when combined with BPaL or RHZE. However, addition of CC214-2 to BPaL and RHZE was associated with significantly fewer relapses in C3HeB/FeJ compared to addition of rapamycin and, in RHZE-treated mice, resulted in fewer relapses compared to RHZE alone. Therefore, CC214-2 and related mTOR kinase inhibitors may be more effective candidates for HDT than rapamycin analogs and may have the potential to shorten the duration of TB treatment.

## Introduction

Despite recent advances in drug development for tuberculosis (TB), first-line treatment for drug-susceptible TB still consists of an extended regimen comprising four drugs: isoniazid, rifampin, pyrazinamide and ethambutol for 2 months, followed by isoniazid and rifampin for another 4 months (RHZE). Treatments for multidrug- and extensively drug-resistant (MDR/XDR) TB are longer and more complex, although the novel short-course regimen of bedaquiline, pretomanid and linezolid (BPaL) recently was approved to treat XDR-TB and MDR-TB patients who lack other options for treatment (1). Efforts to develop more effective and shorter-course therapies for TB have included a focus on host-directed therapy (HDT) (2). The goal of HDT is to modulate the endogenous host response to infection, thereby improving the host’s immune defenses against the pathogen, reducing the duration of antibacterial therapy and/or limiting the amount of lung damage sustained. Given the delicate balance between *Mycobacterium tuberculosis* and the immune response of its host, HDT poses an attractive strategy and has already shown promise by demonstrating reduced lung pathology in mouse and rabbit models of pulmonary TB (3, 4). As an added benefit, HDT approaches should not be adversely affected by antimicrobial drug resistance in the infecting *Mycobacterium tuberculosis* strain.

Autophagy and its degradation pathway has been exploited for therapeutic modulation in various disease states such as metabolic conditions, neurodegenerative diseases, cancers, and infectious diseases (5). Autophagy is an intracellular homeostatic process that removes damaged cellular components and organelles during cellular stress via lysosomal degradation. It is part of the innate immune response involved in eliminating intracellular pathogens (i.e., xenophagy), including *M. tuberculosis* (6, 7). Additionally, it is involved in adaptive immunity by facilitating antigen presentation, which eventually leads to granuloma formation (8). As such, induction of autophagy has become a target of choice for adjunctive HDT in TB (9, 10).

One of the cellular signaling pathways that controls autophagy is regulated by mammalian target of rapamycin (mTOR) (7). While it has been shown that the hypoxic environment in some granulomas inhibits mTOR, inducing autophagy (11), *M. tuberculosis* impedes the host cells’ ability to complete autophagy via modulation of mTOR (12). Therefore, utilizing an mTOR inhibitor could counteract the effect of *M. tuberculosis* infection on mTOR and provide a novel HDT for TB. Although rapamycin, a well-known immunosuppressive drug and commonly used anti-cancer treatment, inhibits mTOR and is an inducer of autophagy, it would have limited utility in TB due to its variable oral absorption and metabolism by CYP3A4, which is induced by the anti-TB drug rifampin (13). The rapamycin analog everolimus has improved bioavailability and recently was evaluated as an adjunct to the standard-of-care chemotherapy regimen in a clinical trial (#NCT02968927). Preliminary interim analyses suggested reduced inflammation and trends toward improved lung function and accelerated culture conversion with addition of everolimus (14). These results suggest that chemical modulation of mTOR and autophagy has potential as a novel HDT approach for TB treatment.

mTOR is a serine/threonine kinase and exists within two distinct multiprotein complexes, mTOR complex-1 (mTORC1) and mTOR complex-2 (mTORC2) (15). Rapamycin and analogs such as everolimus are allosteric inhibitors that target only the mTORC1 complex (16). Based on the modest efficacy seen with rapamycin monotherapy, it has been hypothesized that mTOR kinase inhibitors capable of blocking both mTORC1 and mTORC2 signaling could have expanded therapeutic potential (13) (17). CC214-2, represents a novel series of 4,6-disubstituted-3,4-dihydropyrazino[2,3-b]pyrazin-2(1H)-one selective mTOR kinase inhibitors (18, 19). It is potent, orally bioavailable, directly inhibits both mTORC1 (pS6) and mTORC2 (pAktS473) signaling *in vitro* and in mice and has demonstrated induction of autophagy in U87EGFRvIII xenografts *in vivo* (16, 20). In a mouse model, CC214-2 significantly inhibited PC3 prostate xenograft tumor growth in a dose and schedule-dependent manner at doses up to 100 mg/kg (21). With established pharmacokinetics and pharmacodynamics (PK/PD) parameters, CC214-2 was a viable mTOR inhibitor candidate for TB HDT evaluation.

We hypothesized that the addition of CC214-2 would increase the bactericidal and sterilizing activity of combination chemotherapy for TB in a manner superior to rapamycin. In the present study, we compared the effects of rapamycin and CC214-2, as monotherapies or when added to RHZE or BPaL in chronic low-dose aerosol infection models of TB in BALB/c and C3HeB/FeJ mice. Rapamycin alone promoted more *M. tuberculosis* multiplication (in both mouse strains) and death (in C3HeB/FeJ mice) compared to no treatment, whereas CC214-2 tended to reduce lung bacterial burden. Most importantly, addition of CC214-2 to RHZE significantly reduced the proportion of relapses among C3HeB/FeJ mice, a model recently shown to recapitulate host transcriptional signatures observed in human TB. On the other hand, addition of rapamycin did not improve the efficacy of RHZE in C3HeB/FeJ mice and reduced its efficacy in BALB/c mice. Rapamycin also reduced the efficacy of BPaL in both mouse strains. These results suggest that the mTOR kinase inhibitor CC214-2 may be a more effective and safer mTOR inhibitor candidate for HDT than rapamycin and its analogs and that CC214-2 may have the potential to improve treatment outcomes and perhaps shorten the duration of TB treatment when combined with RHZE.

## Results

### *In vitro* activity of mTOR inhibitors against *M. tuberculosis*

To rule out direct antimicrobial activity of CC214-2, minimum inhibitory concentrations (MICs) of CC214-2 were determined against *M. tuberculosis* HN878 using the broth macrodilution method. The MIC of CC214-2 was >40 μg/mL. Rapamycin was also tested and, consistent with previous observations (22), it inhibited growth of *M. tuberculosis*, but only at a high concentration (MIC = 40 μg/mL).

### CC214-2 pharmacokinetics and dose selection

CC214-2 has demonstrated exposures after oral dosing suitable for evaluation of mTOR inhibition and efficacy *in vivo*. The single-dose plasma PK parameters for CC214-2 doses bracketing the selected 30 mg/kg dose are shown in Table S1. In previous studies, CC214-2 demonstrated significant inhibition of both mTORC1 (pS6) and mTORC2 (pAktS473) for at least 8 and 24 h when dosed at 30 and 100 mg/kg, respectively (21). Furthermore, dosing at 25 and 50 mg/kg once a day in a 21-day PC3 tumor xenograft model resulted in tumor volume reductions of 63% and 84%, respectively, as compared to vehicle control (20) (21). Based on these previous PK/PD and efficacy studies, a dose of 30 mg/kg was selected to best balance predicted long-term tolerability and provide sufficient exposure and target engagement to evaluate the potential of dual mTORC1/mTORC2 inhibition as HDT in murine models of TB.

### Differential effects of mTOR inhibitors on the control of chronic *M. tuberculosis* infection

Schemes for the efficacy experiments in BALB/c and C3HeB/FeJ mice are shown in Tables 3 and 4, respectively. Mean CFU count results for BALB/c and C3HeB/FeJ mice are shown in Tables S5 and S6, respectively.

To determine the effects of the two different classes of mTOR inhibitor against active TB infection, we assessed mouse survival and viable *M. tuberculosis* colony-forming unit (CFU) counts in the lungs before and during monotherapy with rapamycin or CC214-2 in chronic infection models using BALB/c and C3HeB/FeJ mice. As expected, untreated BALB/c mice experienced stable lung infection for 15 weeks post-infection (i.e., up to Week 8 of the treatment period) (Fig. 1A). Compared to these untreated controls, mice treated with rapamycin alone experienced a loss of bacterial containment, resulting in a marked increase in *M. tuberculosis* CFU at Week 8 (p<0.0001), whereas CC214-2 did not adversely affect control of the infection or survival. Unlike BALB/c mice, C3HeB/FeJ mice may develop caseating lung lesions and fail to control low-dose aerosol infections. As shown in Fig. 2, >50% of untreated C3HeB/FeJ mice succumbed to the infection (median survival time, 51 days from Day 0 [99 days post-infection]). Remarkably, all mice treated with rapamycin alone died within six weeks of treatment initiation (median survival time, 24 days from Day 0), which was significantly worse than no treatment (p=0.0195); and among rapamycin-treated mice surviving to Week 4, the lung CFU counts were higher than those in untreated controls (Fig. 3A). In contrast, treatment with CC214-2 alone resulted in only 25% mortality (p=0.0107 vs. rapamycin alone) that occurred only early in the treatment period, but was not statistically significantly different from the survival of untreated mice up to the end of the observation period. CC214-2 treatment reduced CFU counts compared to untreated C3HeB/FeJ controls surviving to Week 8, although the difference was not statistically significant.

**Fig. 1.**
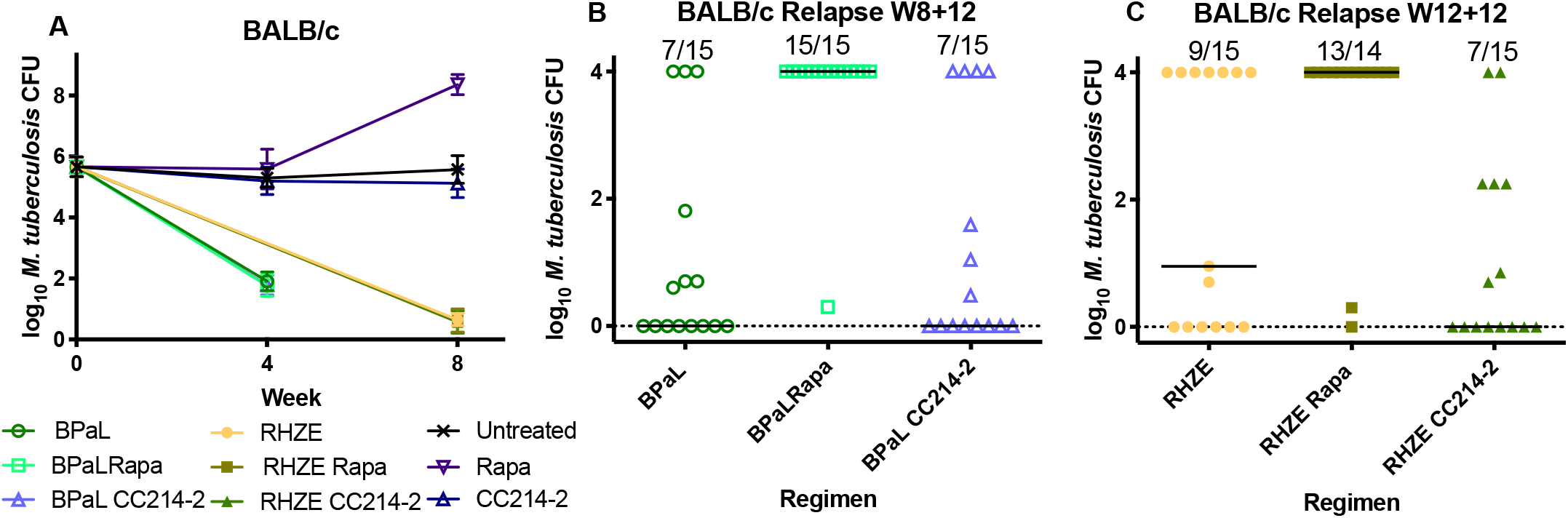
Lung CFU counts in BALB/c mice after treatment. A) Mice treated for one month with BPaL ± an mTOR inhibitor, CC214-2 (CC214), or an mTOR inhibitor alone and for two months with the RHZE regimen ± an mTOR inhibitor. B) Relapse assessment after 8 weeks of treatment with BPaL ± an mTOR inhibitor. C) Relapse assessment after 12 weeks of treatment with RHZE ± an mTOR inhibitor.

**Fig. 2.**
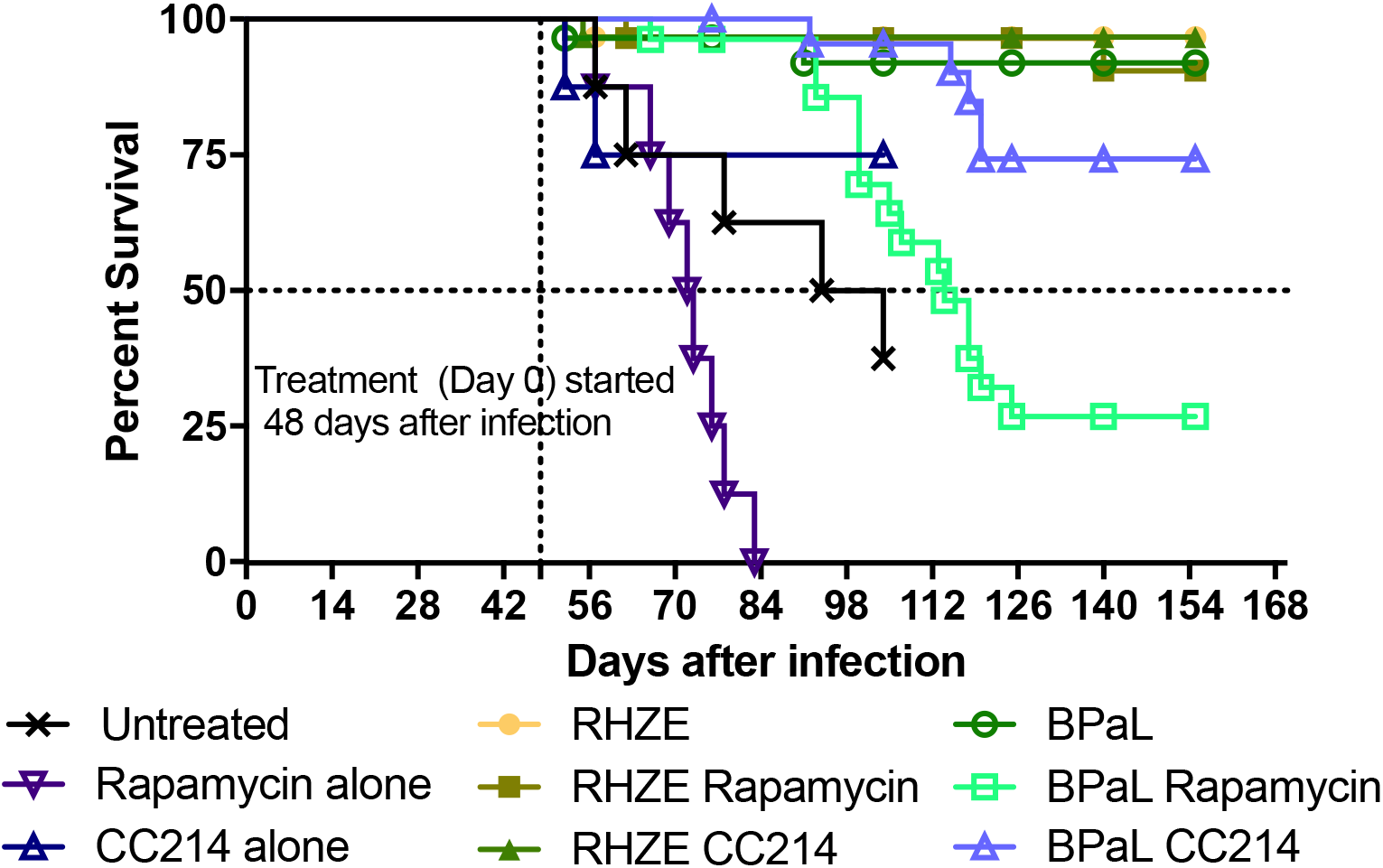
Survival analysis: In C3HeB/FeJ mice, treatment with Rapa alone accelerated weight loss and mortality compared to untreated mice whereas CC214-2 showed a trend towards preserved weight and survival up to the week 8 endpoint. Addition of Rapa to BPaL was also associated with accelerated mortality relative to BPaL alone. Addition of CC214-2 (CC214) to BPaL delayed and fewer deaths compared to Rapa. An adverse effect on mortality or relapse was not observed when Rapa was added to RHZE in C3HeB/FeJ mice.

**Fig. 3.**
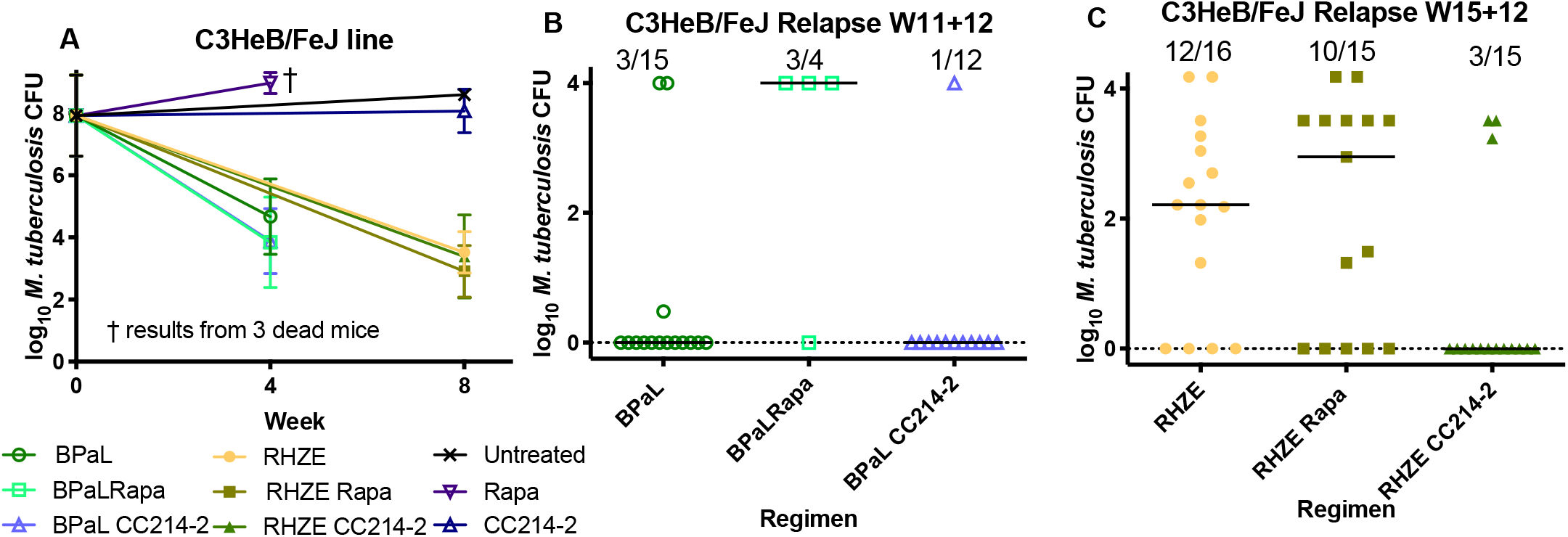
Lung CFU counts in C3HeB/FeJ after treatment. A) Mice treated for one month with the BPaL regimen ± an mTOR inhibitor, CC214-2 (CC214), or rapamycin alone and for two months with the RHZE regimen ± an mTOR inhibitor. No mice treated with rapamycin alone survived to the month 2 timepoint. B) Relapse assessment after 11 weeks of treatment with BPaL ± an mTOR inhibitor. C) Relapse assessment after 15 weeks of treatment with RHZE ± an mTOR inhibitor. NT=not tested

### Differential effects of mTOR inhibitors on the response to combination chemotherapy

To determine the contribution of adjunctive therapy with each mTOR inhibitor to the efficacy of the RHZE and BPaL regimens, BALB/c and C3HeB/FeJ were treated with each regimen alone or in combination with rapamycin or CC214-2. The bactericidal activity of the BPaL and RHZE regimens over 4 and 8 weeks produced the expected reductions of Mtb CFU in both mouse strains (Figs. 1A and 3A) and prevented mortality in C3HeB/FeJ mice (Fig. 2). Neither CC214-2 nor rapamycin altered the lung CFU counts when added to either regimen in either mouse strain. However, in C3HeB/FeJ mice, the addition of rapamycin to BPaL reduced survival compared to BPaL alone (p<0.0001) and BPaL plus CC214-2 (p=0.0008).

The effects of adjunctive mTOR inhibitors on the curative efficacy of the RHZE and BPaL regimens were measured by the proportions of mice with culture-positive lungs (relapse) and the CFU counts in those lungs assessed 12 weeks after completing abbreviated treatment durations. Owing to the superior efficacy of BPaL over RHZE and the typically longer treatment time required for cure in C3HeB/FeJ mice compared to BALB/c mice, BPaL treatment duration was 8 and 11 weeks and RHZE treatment duration was 12 and 15 weeks in BALB/c and C3HeB/FeJ mice, respectively. Treatment with RHZE produced the expected proportions of relapses in both mouse strains. Most importantly, the addition of CC214-2 significantly reduced the proportion of C3HeB/FeJ mice that relapsed (p=0.0038) (Fig. 3C). In contrast, the addition of rapamycin to RHZE did not significantly affect the proportion of mice relapsing, and was less effective than the addition of CC214-2 (p=0.0253). A similar trend was observed in BALB/c mice treated with RHZE with or without CC214-2, although the reductions in the relapse proportion and CFU counts in relapsing mice (Fig. 2C) observed with addition of CC214-2 did not meet statistical significance when compared to RHZE only. However, rapamycin had a deleterious effect on relapse in BALB/c mice (p=0.0142 vs. RHZE only).

Addition of CC214-2 to the novel BPaL regimen resulted in numerically fewer relapses in C3HeB/FeJ, but not BALB/c, mice (Figs. 3B and 2B). This difference in C3HeB/FeJ mice was not statistically significant, but the small number of relapses in these groups could have obscured a larger effect. In stark contrast, the addition of rapamycin to BPaL resulted in significantly greater mortality in C3HeB/FeJ mice (Fig. 2), as well as a higher proportion of mice relapsing compared to BPaL only in both C3HeB/FeJ (p=0.0709) and BALB/c mice (p=0.0022). The small number of assessable C3HeB/FeJ mice treated with BPaL plus rapamycin likely limited the power to detect a statistically significant difference. Still, addition of rapamycin was associated with significantly more relapses than the addition of CC214-2 in both C3HeB/FeJ (p=0.0253) and BALB/c mice (p=0.0022).

### Quantitative histopathology analysis

Overall areas of consolidation and, especially, necrosis tended to be greater in C3HeB/FeJ mice, regardless of treatment with RHZE and/or mTOR inhibitors (Figure S1). Among BALB/c mice, treatment with rapamycin was associated with a trend towards higher scores for consolidation and necrosis relative to no treatment or treatment with CC214-2. Among the sampled C3HeB/FeJ mice, there was no apparent effect of treatment with CC214-2 compared to no treatment, but this comparison may be biased by the loss of more severely affected untreated mice to mortality prior to the Week 8 sampling time point. Similarly, the impact of rapamycin monotherapy in C3HeB/FeJ mice could not be assessed due to the accelerated mortality in that group. Trends towards less necrosis and consolidation were observed when CC214-2 and rapamycin were added to RHZE treatment in C3HeB/FeJ mice, but the small sample numbers precluded statistical analyses or firm conclusions to be drawn. BPaL-treated groups were not assessed.

## Discussion

The most important finding of our study is that the addition of the novel mTOR kinase inhibitor CC214-2 to the first-line RHZE regimen significantly reduced the proportion of C3HeB/FeJ mice relapsing after an abbreviated course of treatment. This result indicates the potential of mTOR kinase inhibition targeting both mTORC1 and mTORC2 to improve TB treatment outcomes and possibly shorten treatment duration in drug-susceptible TB, particularly given recent evidence that this mouse model results in host transcriptional signatures reflective of those seen in TB patients (23). Remarkably, rapamycin, which only partially inhibits mTORC1 signaling through a different mechanism of action, had no evident benefit when added to RHZE. Interestingly, the addition of CC214-2 to the novel BPaL regimen had no significant effect on efficacy, while the addition of rapamycin to BPaL was clearly detrimental, significantly reducing survival in C3HeB/FeJ mice and increasing the proportions of both C3HeB/FeJ and BALB/c mice relapsing after treatment. The discrepancy between the therapeutic effects of the two mTOR inhibitors was further evident in the marked worsening of the infection caused by rapamycin monotherapy compared to an opposing trend toward reduced bacterial load observed with CC214-2 monotherapy in both mouse strains.

The widely discrepant effects observed between the mTOR inhibitors evaluated here may be attributable to the properties and selectivity of the mTOR inhibitors. Although rapamycin and especially analogs such as everolimus have been explored (7, 12, 14, 24, 25) for use as adjunctive HDT for TB, to our knowledge, they have not been studied previously with combination chemotherapy in a mouse model of TB. Due to their effects on the AKT/mTOR pathway and autophagy, they only partially inhibit mTORC1 and do not inhibit mTORC2. The inability of rapamycin to completely block mTORC1-mediated signaling events may be explained by the presence of several feedback loops, and the upregulation of compensatory pathways. There are several feedback loops involved in the cell survival responses, and it has been shown that rapamycin modulates a feedback loop resulting in mTORC2-mediated AKT activation (13). In contrast, inhibitors of mTOR kinase, such as CC214-2, prevent this feedback loop-mediated activation through direct inhibition of pAKT. Additionally, rapamycin does not significantly affect the mTORC1-mediated process of autophagy in that, although rapamycin can effectively activate autophagy in yeast, this induction is limited in mammalian cells (26). In contrast to rapamycin and the “rapalogs”, CC214-2 blocks both mTORC1 and mTORC2 signaling pathways and has demonstrated activation of autophagy in U87EGFRvIII xenografts *in vivo* (16). Taking these mechanistic differences into account, our results strongly suggest the superior therapeutic potential of combined interruption of both mTORC1 and mTORC2 signaling by an mTOR kinase inhibitor as adjunctive HDT for TB, as opposed to the partial inhibition of mTORC1 observed with rapamycin and its analogs. Considering the modest benefits recently observed with addition of everolimus to a regimen of rifabutin plus HZE in a clinical study (14), the therapeutic advantages of CC214-2 observed here suggest that an mTOR kinase inhibitor that more effectively targets both mTORC1 and mTORC2 may deliver superior outcomes in the clinical setting.

Rapamycin’s poor solubility, metabolism by CYP 3A4 and efflux by P-glycoprotein (P-gp) complicate its use as adjunctive HDT for TB. Everolimus, while more soluble and orally bioavailable, is also a substrate of CYP 3A4 and P-gp, and its metabolism is induced by rifampin, the cornerstone drug in the RHZE regimen (27). Hence, the requirement to substitute rifabutin for rifampin when everolimus was evaluated as adjunctive HDT in the aforementioned clinical trial (14). CC214-2 has superior physicochemical and pharmacokinetic properties, and has demonstrated efficacy *in vivo* across multiple solid tumor models when dosed orally. Neither CC214-2 nor the more clinically advanced drug in this class, CC-223, is a substrate, inhibitor or inducer of CYP 3A4 or P-gp, making them more suitable for combination chemotherapy of TB.

Several other findings of the present study warrant further discussion. The benefit of adding CC214-2 to RHZE was more evident in C3HeB/FeJ mice than in BALB/c mice. Despite its relatively recent adoption as a pre-clinical model of TB, the C3HeB/FeJ mouse infection model is now better validated as a model of TB immunopathogenesis than more commonly used mouse strains such as C57Bl/6 and BALB/c mice. Rigorous transcriptomic analyses of peripheral blood and lung samples from infected mice recently indicated a striking resemblance between transcriptional signatures associated with susceptibility to TB disease in C3HeB/FeJ mice and in active human TB (23, 28), including upregulation of type I interferon signaling and neutrophil recruitment and down-regulation of B lymphocyte, natural killer cell and T cell effector responses. These signatures in C3HeB/FeJ mice were clearly more representative of human TB transcriptional signatures than those observed in more resistant C57Bl/6 and BALB/c mice.

Moreover, infection of C3HeB/FeJ mice with the HN878 strain *M. tuberculosis* used in the current study also produced blood and lung transcriptional signatures more similar to human TB signatures than infection with the more commonly used H37Rv strain (23). Thus, the C3HeB/FeJ mouse model used here can be considered highly representative of the host response in human TB disease, and therefore the ideal murine model for investigating HDT candidates. Use of this model increases the probability that the significant benefit of CC214-2 in combination with RHZE observed in this strain will translate to human TB.

Another interesting finding of the present study is the greater benefit observed when CC214-2 was added to the RHZE regimen compared to its addition to the BPaL regimen. Although it is possible that the low number of relapses observed with BPaL treatment, with or without adjunctive CC214-2 treatment, obscured a larger effect of CC214-2 that would have been evident if the duration of treatment had been shorter, it is also possible that the composition of the antimicrobial regimen influenced the magnitude of the CC214-2 effect. An intriguing hypothesis is that mTOR inhibition augments the effects of some TB drugs more than others. Indeed, recent studies suggest that bedaquiline treatment triggers multiple antimicrobial defense mechanisms in macrophages, including lysosomal activation, phagosome-lysosome fusion and autophagy (29, 30). Linezolid also induces autophagy in mammalian cells, apparently through mitochondrial stress and dysfunction (29, 31). Therefore, it is conceivable that the degree to which BPaL itself induces autophagy and other protective antimicrobial mechanisms negates any additional contribution from a direct mTOR kinase inhibitor. In contrast, rifampin, isoniazid and pyrazinamide did not significantly increase lysosomal activation or autolysosome formation to the extent that bedaquiline and linezolid did (29, 30). Although isoniazid and pyrazinamide reportedly activated autophagy in *M. tuberculosis*-infected murine macrophages in one study (6), isoniazid did not increase autophagy in human cells in other studies (29, 30). Furthermore, pyrazinamide’s antibacterial activity is pH-dependent and thus enhanced in acidified phagolysosomes. Increased delivery of *M. tuberculosis* to such phagolysosomes mediated through pharmacologic induction of autophagy may be an additional means by which mTOR inhibition could augment pyrazinamide efficacy. Consistent with this, a recent study using an *in vitro* granuloma model found that everolimus augmented pyrazinamide activity to a more significant extent than its effects on isoniazid activity (24). While we cannot exclude the possibility that co-administration of CC214-2 reduced the exposure to one or more components of the BPaL regimen through an adverse drug-drug interaction, this is unlikely because CC214-2 is not a substrate or inducer of CYP 3A4 or P-gp and was administered separately in time from the antimicrobials to avoid interference with absorption. Therefore, in the context of prior studies, our results suggest that the adjunctive benefits of dual mTORC1/mTORC2 inhibition may be regimen-specific and related to the extent to which the anti-TB drugs themselves affect autophagy and other macrophage defense mechanisms.

A limitation of the present study is the use of only one dose of each mTOR inhibitor. In addition to promoting xenophagy, rapamycin and CC214-2 have potential dose-dependent immunosuppressive effects that could tip the balance in favor of the pathogen and negate adjunctive therapeutic benefits. We used a rapamycin dose that provided analogous efficacy results compared to the 30 mg/kg daily dose of CC214-2 in mouse tumor xenograft models. However, this dose was an order of magnitude higher than the dose used in a study showing enhancement of BCG efficacy against *M. tuberculosis* challenge (75 μg/kg daily) (32).

In conclusion, we have demonstrated that the addition of the mTOR kinase inhibitor CC214-2 significantly increased the sterilizing efficacy of the first-line RHZE regimen in a validated C3HeB/FeJ mouse model of TB. While recently reported clinical trial outcomes suggest that the rapamycin analog everolimus also increases the efficacy of rifabutin plus HZE in TB patients, the superior efficacy of CC214-2 over rapamycin in our study suggests that CC214-2 and other drugs that inhibit both mTORC1 and mTORC2 activity and bypass feedback activation offer greater potential as HDT for TB.

## Materials and Methods

### Bacterial strain

The virulent *M. tuberculosis* HN878 strain, belonging to the Beijing subfamily, was used for infection. The strain was mouse passaged and aliquoted in frozen stocks. The bacteria were grown in Middlebrook 7H9 liquid media supplemented with 10% oleic acid-albumin-dextrose-catalase (OADC) without Tween 80. After growing for five days the culture was de-clumped by passing through a 25 g needle before infection.

### Determination of MICs against *M. tuberculosis* HN878

MICs of rapamycin and CC214-2 were determined using the broth macrodilution method in complete 7H9 media without Tween 80. The drugs were serially diluted 2-fold to obtain the final concentration range from 80 to 1.25 μg/mL. MIC was defined as the lowest concentration to prevent visible growth after 14 days of incubation.

### Aerosol infection of mice with *M. tuberculosis*

All animal procedures were approved by the Animal Care and Use Committee of Johns Hopkins University. Six-week-old female BALB/c mice were purchased from Charles River Laboratories (Wilmington, MA). Eight-to ten-week-old female C3HeB/FeJ mice were purchased from the Jackson Laboratory (Bar Harbor, ME). Mice were infected via aerosol using the GlasCol Inhalation Exposure System. Both BALB/c and C3HeB/FeJ mice were infected using a culture of *M. tuberculosis* HN878 prepared as described above and diluted 1:400 prior to infection. Four mice from each strain were sacrificed and lung homogenates were plated on 7H11 agar plates supplemented with 10% OADC, to estimate the number of implanted CFUs. Seven weeks after infection, C3HeB/FeJ mice were block-randomized into treatment groups according to body weight and aerosol infection run. BALB/c mice were block-randomized by aerosol infection run only. Five BALB/c mice and eight C3HeB/FeJ mice were sacrificed, and lung homogenates were plated to enumerate CFU at the initiation of treatment (Day 0).

### Chemotherapy

The daily doses of drugs (in mg/kg body weight) were 10 (isoniazid, H), 10 (rifampin, R), 100 (ethambutol, E), 150 (pyrazinamide, Z) for the RHZE regimen; 25 (bedaquiline, B), 100 (pretomanid, Pa) 100 (linezolid, L) for the BPaL regimen; 4 (rapamycin, Rapa) and 30 (CC214-2, CC214)(20). R, H, E and Z were prepared in deionized water, B was prepared in acidified HPCD (acidified 20% hydroxypropyl-β-cyclodextrin)solution, Pa in CM2 formulation, L in 0.5% methyl cellulose (33, 34), CC214-2 in 0.5% CMC and 0.25% Tween 80, and rapamycin in 45%PEG 400, 53% saline and 2% ethanol. All drugs were administered orally by gavage 5 days per week except rapamycin, which was administered by intraperitoneal injections 3 days per week (i.e., Mon-Wed-Fri).

### Assessment of treatment efficacy

Treatment efficacy was assessed by evaluating lung CFU counts after 4 or 8 weeks of treatment and the proportions of mice culture-positive 12 weeks after completing different durations of treatment. Lung CFU counts during treatment were determined from a higher number of C3HeB/FeJ mice (n=8) compared to BALB/c mice (n=5) due to the more heterogenous disease progression characteristics of the former strain. Lung homogenates from both groups were serially diluted and plated on selective Middlebrook 7H11 media containing 0.4% activated charcoal to reduce the effects of bedaquiline carryover. CFU counts were determined after 6 weeks of incubation at 37° C. The entire lung homogenates of the relapse cohorts were plated on five plates to reduce the lower limit of CFU detection to 1 when determining the proportion of mice with culture-positive relapse.

### Monitoring clinical signs and recording death of mice

Four weeks (28 days) after infection a baseline body weight of the C3HeB/FeJ mice was assessed. The body weights of the mice were then monitored every week during pretreatment followed by every 2 weeks until completion of the experiment. Physical signs of drug toxicity were recorded. All dead mice were recorded for survival analysis.

### Statistical analysis

CFU counts (x) were log-transformed (as x + 1) before analysis, and group means were compared by one-way analysis of variance with Dunnett’s post-test to control for multiple comparisons. Group relapse proportions were compared using Fisher’s exact test, adjusting for multiple comparisons. The Mann-Whitney test was used to test for significance on non-normally distributed CFU data from C3HeB/FeJ mice. Kaplan-Meier curves and Mantel-Cox logrank tests were used for survival analysis. GraphPad Prism version 6 (GraphPad, San Diego, CA) was used for all analyses. Use of 15 mice per group for relapse assessment provides approximately 80% power to detect 40 percentage point differences in the relapse rate, after setting alpha at 0.01 to adjust for up to 5 simultaneous two-sided comparisons. Smaller differences may not be meaningful in terms of shortening the duration of treatment.

### Quantitative analysis of lung histopathology

Lung samples were collected from both BALB/c and C3HeB/FeJ mice for histopathology analysis after 8 weeks of treatment except for the rapamycin alone group in C3HeB/FeJ mice, which had samples collected after 4 weeks of treatment due to the accelerated mortality observed in that group. Two or three lung samples were collected from each group. Lungs were fixed in 4% paraformaldehyde. The paraffin blocks of lung samples were processed to obtain at least three sections per lung spaced 60 μm apart for hematoxylin and eosin staining. Adjacent sections were stained with Ziehl-Neelsen (acid fast staining) and Masson trichrome stains. Quantitative analysis was performed by a reviewer (MEU) blinded to treatment allocation and mouse strain. For each treatment group, a total of six or nine scanned slides (i.e., from 2 or 3 mice) were reviewed. Regions of interest corresponding to the absolute areas of total lung surface, lesion, inflammation, consolidation and necrosis were manually drawn and their area quantified using the open source software Qupath (https://qupath.github.io/).

## Acknowledgements

This research was supported by TB Alliance (Global Alliance for TB Drug Development), which is funded by Australia’s Department of Foreign Affairs and Trade, Bill & Melinda Gates Foundation [OPP1129600], the Foreign, Commonwealth and Development Office (United Kingdom), Germany’s Federal Ministry of Education and Research through KfW, Irish Aid, Netherlands Ministry of Foreign Affairs, and the United States Agency for International Development. We also thank Drs. Peter Schafer and Khisi Mdluli for helpful discussions and Sandeep Tyagi and Dr. Jian Xu for their assistance with drug dosing of mice.

## References

1. Conradie F, Diacon AH, Ngubane N, Howell P, Everitt D, Crook AM, Mendel CM, Egizi E, Moreira J, Timm J, McHugh TD, Wills GH, Bateson A, Hunt R, Van Niekerk C, Li M, Olugbosi M, Spigelman M. 2020. Treatment of Highly Drug-Resistant Pulmonary Tuberculosis. N Engl J Med 382:893–902.

2. Wallis RS, Hafner R. 2015. Advancing host-directed therapy for tuberculosis. Nat Rev Immunol 15:255–63.

3. Subbian S, Peixoto B, O’Brien P, Dartois V, Kaplan G, Tsenova L, Holloway J, Khetani V, Zeldis JB. 2016. Adjunctive Phosphodiesterase-4 Inhibitor Therapy Improves Antibiotic Response to Pulmonary Tuberculosis in a Rabbit Model. EBioMedicine 4:104–14.

4. Subbian S, Koo M-S, Tsenova L, Khetani V, Zeldis JB, Fallows D, Kaplan G. 2016. Pharmacologic inhibition of host phosphodiesterase-4 improves isoniazid-mediated clearance of Mycobacterium tuberculosis. Front Immunol 7:238/1–238/12.

5. Rubinsztein DC, Codogno P, Levine B. 2012. Autophagy modulation as a potential therapeutic target for diverse diseases. Nat Rev Drug Discov 11:709–30.

6. Kim J-J, Lee H-M, Shin D-M, Kim W, Yuk J-M, Jin HS, Lee S-H, Cha G-H, Kim J-M, Lee Z-W, Shin SJ, Yoo H, Park YK, Park JB, Chung J, Yoshimori T, Jo E-K. 2012. Host cell autophagy activated by antibiotics is required for their effective antimycobacterial drug action. Cell Host Microbe 11:457–468.

7. Singh P, Subbian S. 2018. Harnessing the mTOR Pathway for Tuberculosis Treatment. Front Microbiol 9:70.

8. Cerni S, Shafer D, Venketaraman V, To K, Venketaraman V. 2019. Investigating the Role of Everolimus in mTOR Inhibition and Autophagy Promotion as a Potential Host-Directed Therapeutic Target in Mycobacterium tuberculosis Infection. J Clin Med 8.

9. Deretic V. 2014. Autophagy in tuberculosis. Cold Spring Harb Perspect Med 4:a018481.

10. Torfs E, Piller T, Cos P, Cappoen D. 2019. Opportunities for overcoming Mycobacterium tuberculosis drug resistance: emerging mycobacterial targets and host-directed therapy. Int J Mol Sci 20:2868.

11. Huang HY, Lin PY, Huang CP, Huang HY, Wang WC, Chen CY, Chen YK, Wang WC, Chen CY, Chen YK, Wang WC, Chen CY, Chen YK. 2018. The roles of autophagy and hypoxia in human inflammatory periapical lesions. Int Endod J 51 Suppl 2:e125–e145.

12. Lachmandas E, Beigier-Bompadre M, Cheng SC, Kumar V, van Laarhoven A, Wang X, Ammerdorffer A, Boutens L, de Jong D, Kanneganti TD, Gresnigt MS, Ottenhoff TH, Joosten LA, Stienstra R, Wijmenga C, Kaufmann SH, van Crevel R, Netea MG. 2016. Rewiring cellular metabolism via the AKT/mTOR pathway contributes to host defence against Mycobacterium tuberculosis in human and murine cells. Eur J Immunol 46:2574–2586.

13. Li J, Kim SG, Blenis J. 2014. Rapamycin: one drug, many effects. Cell metabolism 19:373–379.

14. Wallis RS, Ginindza S, Beattie T, Arjun N, Likoti M, Edward V, Rassool M, Ahmed K, Fielding K, Ahidjo B, Vangu MdT, Churchyard G. Preliminary Results of an Experimental Medicine Trial of Adjunctive Host-Directed Therapy in Adults with Moderately or Far-Advanced Rifampin-Susceptible Pulmonary Tuberculosis, p A7388. In (ed), American Journal of Respiratory and Critical Care Medicine,

15. Shor B, Gibbons JJ, Abraham RT, Yu K. 2009. Targeting mTOR globally in cancer: thinking beyond rapamycin. Cell Cycle 8:3831–7.

16. Gini B, Zanca C, Guo D, Matsutani T, Masui K, Ikegami S, Yang H, Nathanson D, Villa GR, Shackelford D, Zhu S, Tanaka K, Babic I, Akhavan D, Lin K, Assuncao A, Gu Y, Bonetti B, Mortensen DS, Xu S, Raymon HK, Cavenee WK, Furnari FB, James CD, Kroemer G, Heath JR, Hege K, Chopra R, Cloughesy TF, Mischel PS. 2013. The mTOR Kinase Inhibitors, CC214-1 and CC214-2, Preferentially Block the Growth of EGFRvIII-Activated Glioblastomas. Clin Cancer Res 19:5722–5732.

17. Cloughesy TF, Yoshimoto K, Nghiemphu P, Brown K, Dang J, Zhu S, Hsueh T, Chen Y, Wang W, Youngkin D, Liau L, Martin N, Becker D, Bergsneider M, Lal A, Green R, Oglesby T, Koleto M, Trent J, Horvath S, Mischel PS, Mellinghoff IK, Sawyers CL. 2008. Antitumor activity of rapamycin in a phase I trial for patients with recurrent PTEN-deficient glioblastoma. PLoS Med 5:139–151.

18. Mortensen DS, Perrin-Ninkovic SM, Shevlin G, Zhao J, Packard G, Bahmanyar S, Correa M, Elsner J, Harris R, Lee BGS, Papa P, Parnes JS, Riggs JR, Sapienza J, Tehrani L, Whitefield B, Apuy J, Bisonette RR, Gamez JC, Hickman M, Khambatta G, Leisten J, Peng SX, Richardson SJ, Cathers BE, Canan SS, Moghaddam MF, Raymon HK, Worland P, Narla RK, Fultz KE, Sankar S. 2015. Discovery of Mammalian Target of Rapamycin (mTOR) Kinase Inhibitor CC-223. J Med Chem 58:5323–5333.

19. Mortensen DS, Perrin-Ninkovic SM, Shevlin G, Elsner J, Zhao J, Whitefield B, Tehrani L, Sapienza J, Riggs JR, Parnes JS, Papa P, Packard G, Lee BGS, Harris R, Correa M, Bahmanyar S, Richardson SJ, Peng SX, Leisten J, Khambatta G, Hickman M, Gamez JC, Bisonette RR, Apuy J, Cathers BE, Canan SS, Moghaddam MF, Raymon HK, Worland P, Narla RK, Fultz KE, Sankar S. 2015. Optimization of a Series of Triazole Containing Mammalian Target of Rapamycin (mTOR) Kinase Inhibitors and the Discovery of CC-115. J Med Chem 58:5599–5608.

20. Mortensen DS, Sapienza J, Lee BGS, Perrin-Ninkovic SM, Harris R, Shevlin G, Parnes JS, Whitefield B, Hickman M, Khambatta G, Bisonette RR, Peng S, Gamez JC, Leisten J, Narla RK, Fultz KE, Sankar S. 2013. Use of core modification in the discovery of CC214-2, an orally available, selective inhibitor of mTOR kinase. Bioorg Med Chem Lett 23:1588–1591.

21. Mortensen DS, Sapienza JJ, Perrin-Ninkovic SM, Harris R, Lee BG, Shevlin G, Elsner J, Zhao J, Parnes J, Packard GK, Tehrani L, Papa P, Whitefield B, Riggs JR, Correa M, Bahmanyar S, Hickman M, Khambatta G, Bisonette RR, Fultz KE, Gamez J, Leisten J, Peng S, Narla RK, Sankar S. Use of core modification to identify a new series of mTOR kinase inhibitors and CC214-2, an orally available, selective inhibitor of mTOR kinase, p MEDI-293. In (ed), American Chemical Society,

22. Greenstein RJ, Su L, Shahidi A, Brown WD, Clifford A, Brown ST. 2014. Unanticipated *Mycobacterium tuberculosis* complex culture inhibition by immune modulators, immune suppressants, a growth enhancer, and vitamins A and D: clinical implications. Int J Infect Dis 26:37–43.

23. Moreira-Teixeira L, Tabone O, Graham CM, Singhania A, Stavropoulos E, Redford PS, Chakravarty P, Priestnall SL, Suarez-Bonnet A, Herbert E, Mayer-Barber KD, Sher A, Fonseca KL, Sousa J, Cá B, Verma R, Haldar P, Saraiva M, O’Garra A. 2020. Mouse transcriptome reveals potential signatures of protection and pathogenesis in human tuberculosis. Nat Immunol 21:464–476.

24. Ashley D, Hernandez J, Cao R, To K, Yegiazaryan A, Abrahem R, Nguyen T, Owens J, Lambros M, Subbian S, Venketaraman V. 2020. Antimycobacterial Effects of Everolimus in a Human Granuloma Model. J Clin Med 9:2043.

25. Cerni S, Shafer D, To K, Venketaraman V. 2019. Investigating the role of everolimus in mTOR inhibition and autophagy promotion as a potential host-directed therapeutic target in Mycobacterium tuberculosis infection. J Clin Med 8:232.

26. Thoreen CC, Sabatini DM. 2009. Rapamycin inhibits mTORC1, but not completely. Autophagy 5:725–726.

27. Kovarik JM, Hartmann S, Figueiredo J, Rouilly M, Port A, Rordorf C. 2002. Effect of rifampin on apparent clearance of everolimus. Ann Pharmacother 36:981–5.

28. Moreira-Teixeira L, Stimpson PJ, Stavropoulos E, Hadebe S, Chakravarty P, Ioannou M, Aramburu IV, Herbert E, Priestnall SL, Suarez-Bonnet A, Sousa J, Fonseca KL, Wang Q, Vashakidze S, Rodríguez-Martínez P, Vilaplana C, Saraiva M, Papayannopoulos V, O’Garra A. 2020. Type I IFN exacerbates disease in tuberculosis-susceptible mice by inducing neutrophil-mediated lung inflammation and NETosis. Nat Commun 11:5566.

29. Genestet C, Bernard-Barret F, Hodille E, Ginevra C, Ader F, Goutelle S, Lina G, Dumitrescu O. 2018. Antituberculous drugs modulate bacterial phagolysosome avoidance and autophagy in Mycobacterium tuberculosis-infected macrophages. Tuberculosis (Edinb) 111:67–70.

30. Giraud-Gatineau A, Coya JM, Maure A, Biton A, Thomson M, Bernard EM, Marrec J, Gutierrez MG, Larrouy-Maumus G, Brosch R, Gicquel B, Tailleux L. 2020. The antibiotic bedaquiline activates host macrophage innate immune resistance to bacterial infection. Elife 9:e55692.

31. Abad E, García-Mayea Y, Mir C, Sebastian D, Zorzano A, Potesil D, Zdrahal Z, Lyakhovich A, Lleonart ME. 2019. Common Metabolic Pathways Implicated in Resistance to Chemotherapy Point to a Key Mitochondrial Role in Breast Cancer. Mol Cell Proteomics 18:231–244.

32. Jagannath C, Bakhru P. 2012. Rapamycin-induced enhancement of vaccine efficacy in mice. Methods Mol Biol 821:295–303.

33. Rosenthal IM, Tasneen R, Peloquin CA, Zhang M, Almeida D, Mdluli KE, Karakousis PC, Grosset JH, Nuermberger EL. 2012. Dose-ranging comparison of rifampin and rifapentine in two pathologically distinct murine models of tuberculosis. Antimicrob Agents Chemother 56:4331–40.

34. Tasneen R, Betoudji F, Tyagi S, Li SY, Williams K, Converse PJ, Dartois V, Yang T, Mendel CM, Mdluli KE, Nuermberger EL. 2016. Contribution of Oxazolidinones to the Efficacy of Novel Regimens Containing Bedaquiline and Pretomanid in a Mouse Model of Tuberculosis. Antimicrob Agents Chemother 60:270–7.

